# Disentangling metabolic functions of bacteria in the honey bee gut

**DOI:** 10.1101/157461

**Authors:** Lucie Kešnerová, Ruben A. T. Mars, Kirsten M. Ellegaard, Michaël Troilo, Uwe Sauer, Philipp Engel

## Abstract

It is presently unclear how much individual community members contribute to the overall metabolic output of a gut microbiota. To address this question, we used the honey bee, which harbors a relatively simple and remarkably conserved gut microbiota with striking parallels to the mammalian system and importance for bee health. Using untargeted metabolomics, we profiled metabolic changes in gnotobiotic bees that were colonized with the complete microbiota reconstituted from cultured strains. We then determined the contribution of individual community members in mono-colonized bees, and recapitulated our findings using *in vitro* cultures. Our results show that the honey bee gut microbiota utilizes a wide range of pollen-derived substrates including flavonoids and outer pollen wall components, suggesting a key role for degradation of recalcitrant secondary plant metabolites and pollen digestion. In turn, multiple species were responsible for the accumulation of organic acids and polyphenol degradation products, and a specific gut symbiont, *Bifidobacterium asteroides*, stimulated the production of host hormones known to impact bee development. While we found evidence for cross-feeding interactions, ∼80% of the identified metabolic changes were also observed in mono-colonized bees with *Lactobacilli* being responsible for the largest share of the metabolic output. These results suggest that bacteria in the honey bee gut colonize largely independent metabolic niches, which may be a general characteristic of gut microbiomes. Our study reveals diverse metabolic functions of gut bacteria that are likely to contribute to bee health, and provide fundamental insights into how metabolic functions are partitioned within gut communities.

**Author summary:** Honey bees are important pollinators that harbor a simple gut microbiota with striking parallels to the mammalian system. This makes them relevant models to study gut microbiota functions and its impact on host health. We applied untargeted metabolomics to characterize metabolic changes induced by the gut microbiota, and to characterize contributions of the major community members. We find that the gut microbiota digests recalcitrant substrates derived from the bees’ pollen-diet. Most metabolic changes could be explained by the activity of individual community members suggesting substrate specificity and independent metabolic niches. Our study provides novel insights into the functional understanding of the bee gut microbiota and provides a framework for applying untargeted metabolomics to disentangle metabolic functions of gut bacteria.

## Introduction

Metabolic activities of the microbiota are key for establishing mutualistic interactions in the gut and thereby affect the host in conditions of health and disease. Gut bacteria facilitate the breakdown of refractory or toxic dietary compounds [1-3], produce metabolites that promote host growth and physiology [4-7], and modulate immune functions in the gut [8] and other tissues [9,10]. Moreover, metabolic activity is the basis for energy and biomass production resulting in bacterial growth and the occupation of ecological niches conferring colonization resistance against pathogenic microbes [11]. Substrates of gut bacteria predominantly originate from the diet of the host [2,12], making diet the major modulator of the composition and metabolic activity of the gut microbiota [13,14].

The substantial metabolic potential of the animal gut microbiota has been profiled by the direct sequencing of functional gene content (i.e. shotgun metagenomics) [15-18]. However, it is challenging to predict functional metabolic output from such sequencing data. With recent advances in the coverage and throughput of untargeted screening metabolomics [19-21], it has become feasible to quantify metabolic changes in microbiota or host tissues at large coverage and throughput. Besides identifying metabolites connected to human health and disease [22-30], untargeted screening metabolomics holds considerable promise to unravel metabolic functions of individual microbiota members in animals with divergent dietary preferences. Such mono-colonization studies are however complicated by the highly variable and species-rich composition of most animal microbiota. Thus, gut communities of reduced complexity are valuable models to disentangle metabolic functions of the constituent species.

Like mammals, honey bees harbor a highly specialized gut microbiota. However, the honey bee gut microbiota is surprisingly simple and conserved with seven species (categorized by clustering at 97% 16S rRNA sequence identity) accounting for >90% of the entire gut community in bees sampled across continents [31]. This microbiota is composed of four Proteobacteria (*Gilliamella apicola, Snodgrassella alvi, Frischella perrara, Bartonella apis*), which mostly reside in the ileum, and two Firmicutes (*Lactobacillus* spp. Firm-4 and Firm-5) and one Actinobacterium (*Bifidobacterium asteroides*), which are predominantly found in the rectum. These specific locations suggest that bacteria occupy different metabolic niches in the bee gut and potentially engage in syntrophic interactions [32,33].

The honey bee gut microbiota has marked effects on the host. It promotes host weight gain and hormone signaling under laboratory setting [34] and stimulates the immune system of the host [35,36]. In addition, honey bees are ecologically and economically essential pollinators that have experienced increased mortality in recent years [37,38], which could in part be due to disturbances of their microbiota composition [39-42].

Genomic analyses in combination with *in vitro* cultures have shown that fermentation of sugars and complex carbohydrates (e.g. pectin) [15,32,43,44] into fermentation products is the dominant metabolic activity of the gut microbiota [34]. Lacking however is a detailed understanding of the consumption of diet-derived substrates and how individual community members contribute to the metabolic activities *in vivo.* It is for instance elusive whether analogously to mammals recalcitrant dietary compounds (especially from pollen) are broken down by the microbiota in the hindgut (i.e. large intestine composed of ileum and rectum), while more accessible compounds are reportedly absorbed by the host in the midgut (i.e. small intestine) [45-47].

To profile the metabolic output of the honey bee gut microbiota and its individual members, we employed gnotobiotic bee colonizations and *in vitro* experiments in conjunction with untargeted metabolomics **(Figure 1)**. We first characterized robust metabolic differences between microbiota-free bees and bees colonized with a reconstituted community composed of the seven major bacterial species of the gut microbiota. Subsequently, we analyzed bees colonized with each community member separately to assay their potential contribution to the overall metabolic output of the gut microbiota. Finally, we recapitulated our results *in vitro* using pollen-conditioned medium. Our systematic approach provides unprecedented insights into the metabolic activities of the honey bee gut microbiota and demonstrates the usefulness of metabolomics in combination with gnotobiotic animal models to disentangle functions of individual gut microbiota members.

**Figure 1.**
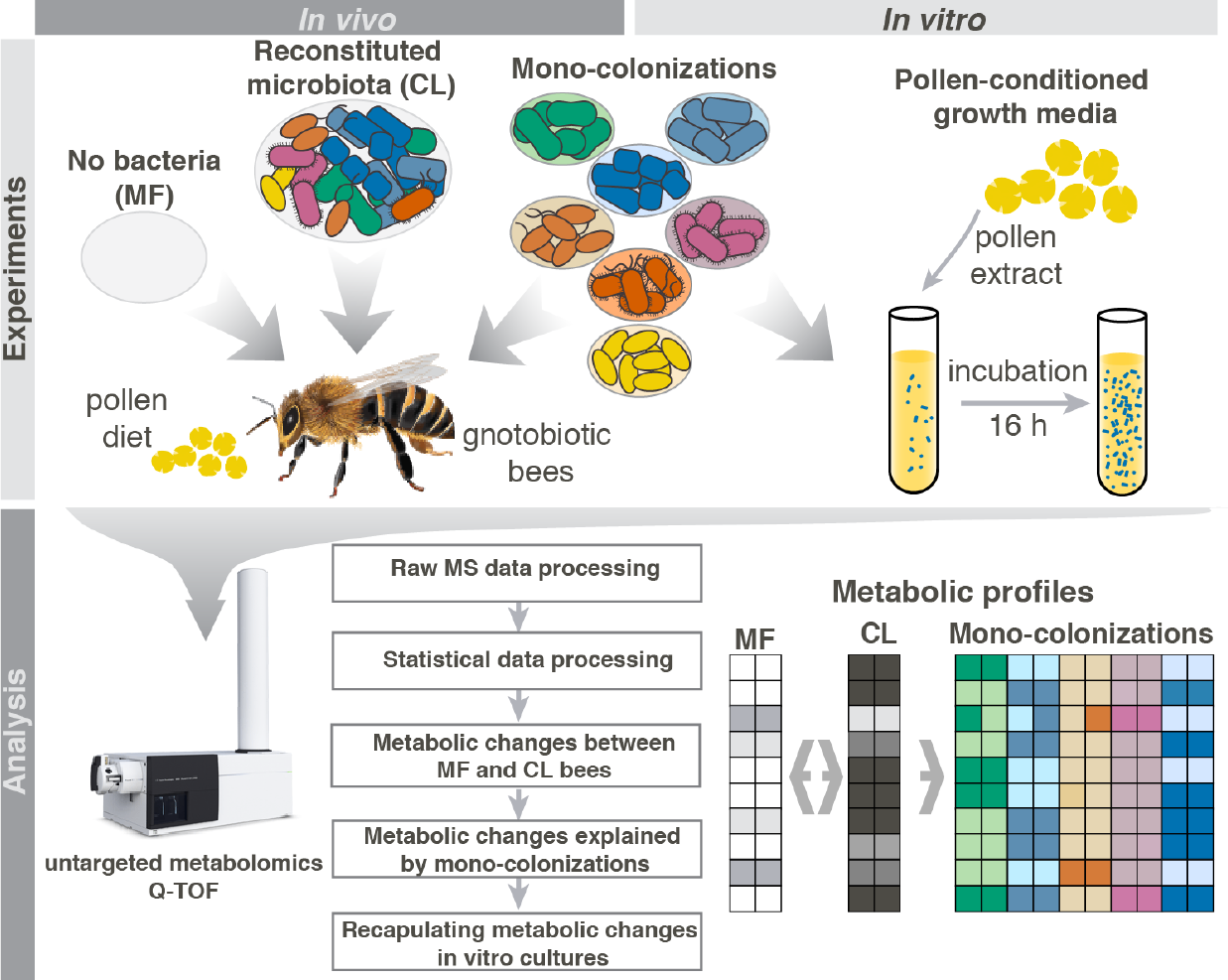
Overview of the experimental setup to characterize metabolic activities of the honey bee gut microbiota. Newly emerged adult bees were either kept microbiota-free, colonized with a reconstituted community of the seven predominant species of the bee gut microbiota (CL), or monocolonized with one of the seven species separately. Bees obtained gamma-sterilized bee pollen as diet. Ten days after colonization honey bee guts were extracted and subjected to untargeted metabolomics to (i) reveal overall metabolomics changes in CL vs MF bees and (ii) to identify which community member could explain these metabolic changes in the gut. As a control, we additionally analyzed 10-day old hive bees that were colonized by the native microbiota under natural conditions in the colony (not shown in this figure). To recapitulate findings *in vitro*, individual community members were cultured in pollen-conditioned medium and metabolic changes in this medium were again profiled using untargeted metabolomics.

## Results and Discussion

### Experimental reconstitution of the honey bee gut microbiota

To characterize the metabolic output of the honey bee gut microbiota, we colonized microbiota-free (MF) bees with selected bacterial strains previously isolated from the bee gut. The reconstituted bacterial community consisted of eleven strains **(Table S1)** covering the seven species described above. We used two strains for *G. apicola* and four strains for Firm-5 in order to cover the extensive genetic diversity within these species [44,48]. Exposure of MF bees to this community resulted in the successful establishment of all seven species with a total of 10^9^ – 10^10^ bacterial cells per gut after 10 days of colonization **(Figure 2, Figure S1A)**. In contrast, non-colonized MF bees had bacterial levels <10^5^ per gut, an observation consistent with previous studies [32,49]. Compared to hive bees of the same age, bacterial abundances of most species were slightly elevated in colonized (CL) bees. However, in both groups the Firm-5 species was consistently the most abundant community member, while *B. apis* colonized at much lower levels. This is in line with recent 16S rRNA gene-based community analysis [31,50,51], and we thus conclude that the selected strains assembled into a structured community resembling the native honey bee gut microbiota. This validates our gnotobiotic bee system as a tool for microbiota reconstitution experiments and enables the study of microbiota functions under controlled laboratory conditions.

**Figure 2.**
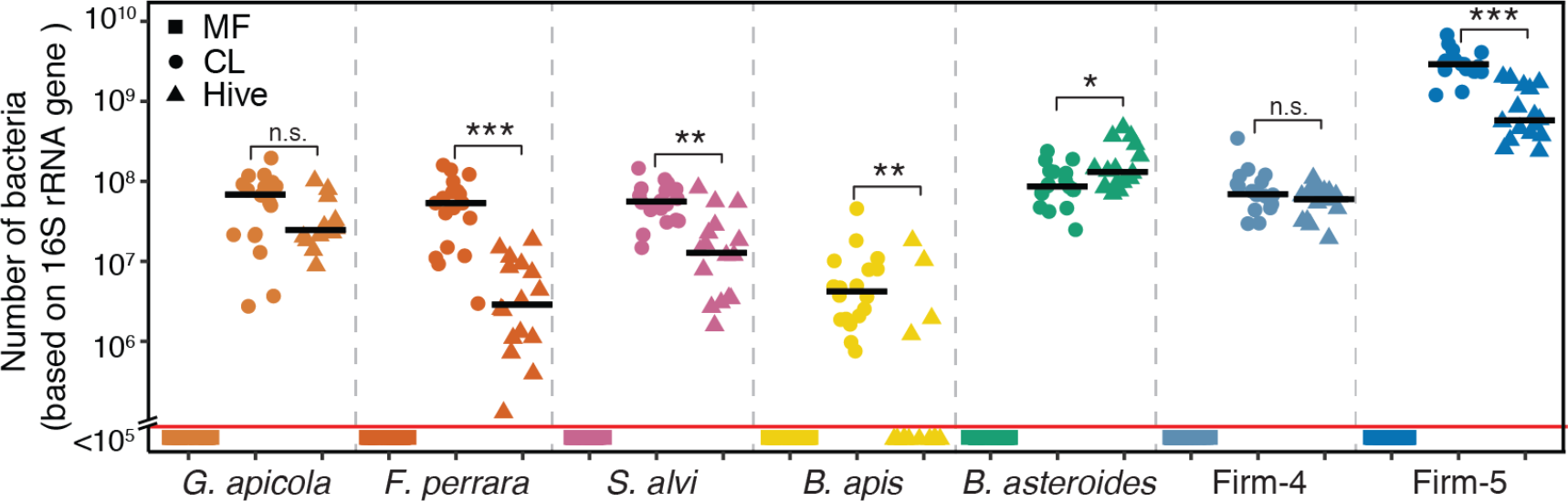
Colonization levels of the seven community members in guts of microbiota-free bees (MF), colonized bees (CL), and hive bees. (A) Bacterial loads in the mid/hindguts of 10-day old MF bees (n= 21), CL bees (n= 18), and hive bees (n = 16) were assessed by quantitative PCR of the 16S rRNA gene using species-specific primer pairs. Black lines show median values. Samples shown below the red line had <10^5^ bacterial cells per gut. Primer characteristics are summarized in Table S5.

### Metabolic changes in the honey bee gut upon colonization with the reconstituted bacterial community

To reveal microbiota-induced metabolome changes in the gut, we dissected the combined mid and hindgut of MF, CL and hive bees and analyzed water-extracted homogenates of these gut samples by untargeted metabolomics [21]. In total, we detected 24,899 mass-to-charge features (ions), 1,079 of which could be annotated by matching their accurate mass to sum formulas of compounds in the full KEGG database **(Dataset S1)**. These 1,079 ions putatively correspond to 3,270 metabolites, since this method cannot separate isobaric compounds **(Supplemental methods)**. For ion changes with multiple annotations, we provided the most likely annotation as based on information from literature and genomic data.

Principal component analysis on the ion intensities revealed that CL and MF bees separate into two distinct clusters suggesting colonization-specific metabolic profiles **(Figure S1B)**. In two independent experiments, a total of 372 ions exhibited significant changes between CL and MF bees (Welch’s t-test, BH adj. *P* ≤0.01). A subset of 240 ions (65%) were more abundant in MF bees suggesting that the cognate metabolites are utilized by the gut microbiota. These ions are hereafter referred to as bacterial substrates. Conversely, 132 ions were more abundant in CL bees and are hereafter referred to as bacterial products indicating that they are produced either by the microbiota or by the host in response to the microbiota. To facilitate the biological interpretation of these multitude metabolic changes, we carried out two analyses. First, we looked whether certain compound classes were overrepresented among the subsets of bacterial substrates and products **(Dataset S2)**. Second, we sorted ion changes based on their ability to explain the difference between the MF and CL metabolome profiles in an Orthogonal Projection of Least Squares Differentiation Analysis (OPLS-DA) [52] **(Figure 3)**.

**Figure 3.**
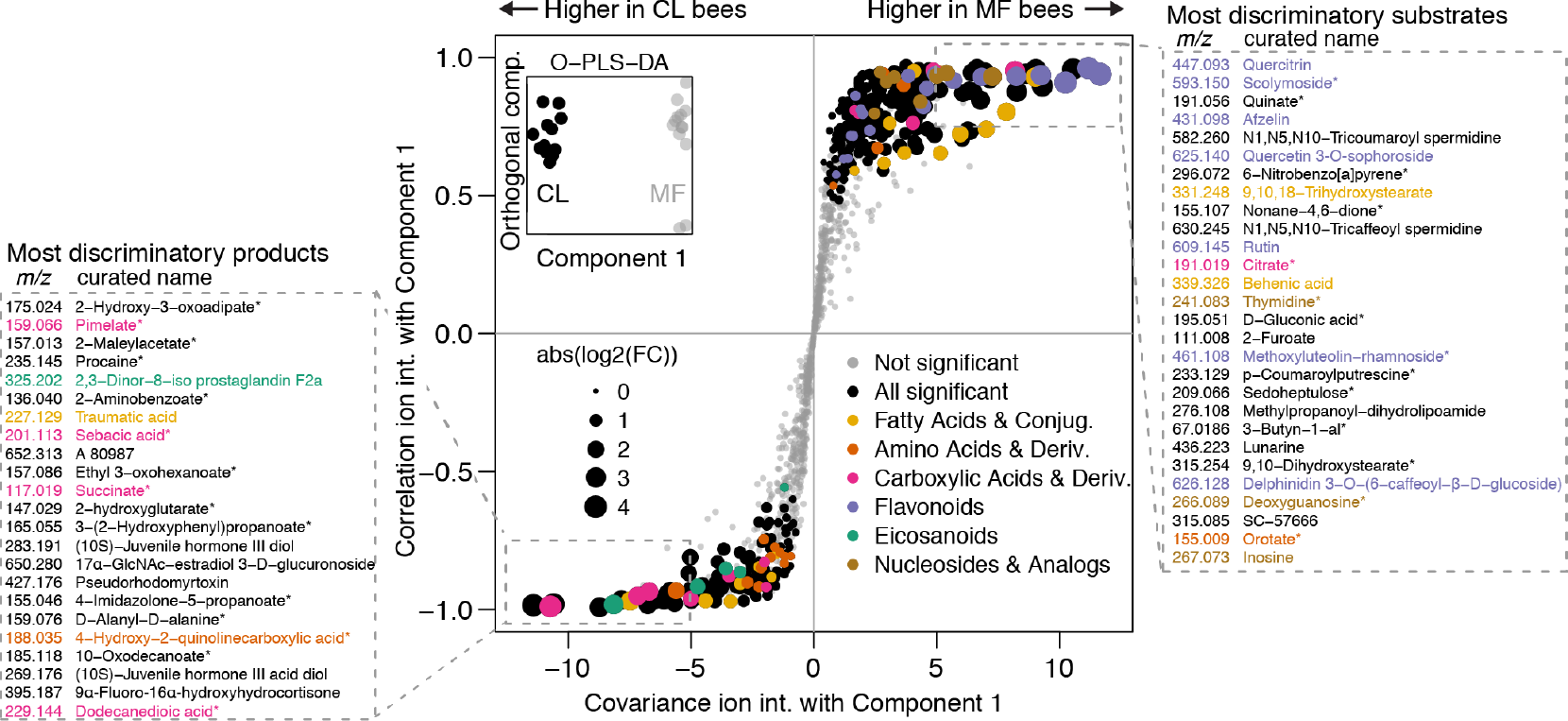
Metabolite changes between microbiota-free (MF) bees and bees colonized with the reconstituted microbiota (CL). Orthogonal Projection of Least Squares-Differentiation Analysis (OPLS-DA) based S-plot of metabolite changes shows ions responsible for MF and CL separation. Inset shows OPLS-DA separation between MF and CL along the component that was used for correlating ion intensities. Experiment 2 data (see **Dataset 1**) was used for this plot, and annotated ions that were not robustly significantly different between MF and CL in both experiments are plotted in grey. Ions with a first annotation belonging to an enriched category are plotted in color (see **Dataset 2**), except for the category *amino acids and derivatives* which did not meet the significance threshold for enrichment but was deemed relevant. The *purine-* and *pyrimidine nucleosides and analogues* categories were combined into *nucleosides and analogs* for coloring only. The boxed areas show the *m/z* [M-H+]^_^of the ion and the first annotation name of the most discriminatory ions, sorted by covariance. Asterisks indicate ions with ambiguous annotations.

### Major bee gut microbiota substrate classes

We first focused on the 240 substrate ions that were more abundant in MF vs CL bees to reveal metabolites potentially utilized by the microbiota. We found three compound classes to be strongly enriched: *flavonoids* (20 of 36 annotated ions, one-sided Fisher’s exact test, *P* <0.001), and both *purine-* and *pyrimidine nucleosides* (in total 8 of 9 annotated ions, both *P* <0.01). Seven flavonoids, three nucleosides, and a nucleoside precursor (orotate, *m/z* 155.009) were also among the 28 substrate ions with most discriminatory power for distinguishing CL vs MF bees as based on OPLS-DA **(Figure 3)**. Other ions among these most discriminatory substrates included two ω-hydroxy acids (*m/z* 315.254 and *m/z* 331.249) and three phenolamides (*m/z* 582.260, *m/z* 630.245, *m/z* 233.130) from the outer pollen wall, as well as quinate (*m/z* 191.056), and citrate (*m/z* 191.019), both of which had previously been predicted to be utilized by certain community members of the honey bee gut microbiota [53,54]. Because they are the most remarkable groups among the identified substrates, nucleosides, flavonoids, and pollen-specific compounds will be discussed in more detail.

#### Nucleosides

Nucleosides are essential biomass components that can also serve as bacterial energy sources. The majority of the honey bee gut microbiota members lack genetic capacity for either purine or pyrimidine biosynthesis [33]. Instead, they seem to rely on salvage pathways and the uptake of nucleosides from the environment, providing a likely explanation for the observed depletion of these compounds in the gut of CL bees **(Figure S2A)**.

#### Flavonoids

Flavonoids are secondary metabolites that can make up 2-4% of the dry weight of pollen [55,56] and this is in concordance with their identification as important diet-derived substrates of the honey bee gut microbiota. Flavonoids typically consist of two phenyl rings and one heterocyclic ring, the so-called aglycone. Glycosylation of this aglycone in diverse positions generates the remarkable flavonoid structural and functional diversity [57]. Among the bacterial substrates we identified several glycosylated flavonoids **(Figure S2B)** that are known to be present in pollen, such as rutin or quercitrin [55]. We unambiguously confirmed the identity of five of these ions as the glycosylated flavonoids afzelin, quercitrin, and rutin using MS/MS fragmentation and spectral similarity calculations **(Supplemental results 1, Figure S3, Dataset S3)**.

Several mammalian gut bacteria can convert flavonoids by either deglycosylation, thus releasing flavonoid aglycones, or C-ring cleavage, resulting in the accumulation of breakdown products of the aromatic backbone [58]. We mined the 132 bacterial products to look for these signatures of flavonoid conversions in the bee gut, and identified several ions annotated as non-glycosylated flavonoids **(Figure S2B)**. However, none of them significantly accumulated in CL vs MF bees, which would have been expected when deglycosylation was the only mechanism of flavonoid conversion. Conversely, we identified four ions among the bacterial products that could result from biodegradation of flavonoid aglycones: two ions annotated as hydroxy‐ and dihydroxyphenylpropionate (*m/z* 165.055, *m/z* 181.050), both of which are known C-ring cleavage products of flavonoids, and two ions annotated as maleylacetate (*m/z* 157.014) and hydroxy-3-oxoadipate (*m/z* 175.025), which are intermediates of aromatic compound degradation pathways [59-62] **(Figure S2B)**. Strikingly, three of these four ions were among the most discriminatory products for CL vs MF bees **(Figure 3)**. Moreover, in a recent metabolomics study by Zheng et al (2017) similar aglycone breakdown products were shown to accumulate in the hindgut of colonized bees **(Supplemental results 2, Dataset S4)**.

#### Pollen wall compounds

The two ions annotated as ω-hydroxy acids (9,10,18‐ trihydroxystearate, *m/z* 315.254, and 9,10-dihydroxystearate, *m/z* 331.249) that were identified among the most discriminatory substrate ions **(Figure 3, Figure S2C)** have been reported to be major constituents of sporopollenin [63], the biochemically inert and heterogeneous biopolymer forming the rigid structure of the pollen exine [64]. Phenolamides such as those utilized by the gut microbiota (N1,N5,N10-tricoumaroyl spermidine, *m/z* 582.260, N1,N5,N10-tricaffeoyl spermidine, *m/z* 630.245, p-coumaroylputrescine, *m/z* 233.130, Figure 3, **Figure S2C)** are also part of the pollen exine as they are deposited on top and into its cavities as part of the pollen coat [65]. Remarkably, the pollen coat is also where most flavonoids are thought to be located in pollen grains [66].

Our findings on the utilization of flavonoids, ω-hydroxy acids and phenolamides thus suggest that the honey bee gut microbiota contributes to the digestion of the rigid outer pollen wall. Easily accessible pollen nutrients (such as amino acids, sugars and vitamins) are likely taken up by the host in the midgut, leaving these more recalcitrant compounds for the microbiota in the hindgut. This is in line with what is know about the biogeography and microbial ecology in the mammalian intestine [67]. Besides being utilized as an energy and carbon source, the conversion of secondary plant metabolites from pollen may have additional benefits for the microbiota and the host. For example, phenolamides and flavonoids both have been reported to exert antimicrobial activities, conceivably because their breakdown products affect the antioxidant potential in the gut which could reduce inflammation and pathogen susceptibility [65,68]. In addition, mammalian flavonoid-metabolizing bacteria have a major impact on the bioavailability of flavonoids, and flavonoids are implicated in modulating weight gain by affecting host signaling [69,70]. This makes it tempting to speculate that flavonoid metabolism in the bee gut contributes to the microbiota‐ dependent weight gain of honey bees that was observed in a previous study [34].

### Products of the gut microbiota include fermentation products and host‐ derived metabolites

We next looked into the 132 ions that were more abundant in CL vs MF bees and thus present possible metabolites produced by the microbiota. Again, we used enrichment analyses and OPLS-DA **(Figure 3)** to prioritize the most important product ions. Three compound classes were to some extent enriched among the bacterial products: *carboxylic acids and derivatives* (7 of 26 detected ions, one sided Fisher’s exact test, *P*<0.03), *fatty acids and derivatives* (7 of 29 detected ions *P* <0.05), and *eicosanoids* (5 of 8 detected ions, *P* <0.01).

#### Fermentation products

Both the *carboxylic acids and derivatives* and *fatty acids and derivatives* categories contain known bacterial fermentation products, several of which accumulated in CL bees (succinate, *m/z* 117.019, pimelate *m/z* 159.066, sebacic acid, *m/z* 201.113, butyrate, *m/z* 87.044, valerate *m/z* 101.060). This is in agreement with previous studies showing that fermentation is the dominant metabolism of bee gut bacteria [33,34]. Three of these fermentation products (succinate, pimelate, sebacic acid) were among the 23 most discriminatory products, which highlights the substantial and consistent accumulation of these compounds in the presence of the microbiota **(Figure 3, Figure S2D)**. Using targeted metabolomics [71] we confirmed the strong accumulation of succinate in the gut of CL bees and determined absolute concentrations of other organic acids in CL and MF bees **(Supplemental results 3, Figure S4, Dataset S5)**. In addition, accumulation of fermentation products is one of the main microbiota-dependent trends found in our study and that of Zheng et al (for details on the comparison see Supplemental results 2).

#### Host-derived metabolites

The third enriched product category (eicosanoids) includes five ions whose masses match to prostaglandins **(Figure S2E)**, which are broadly conserved hormone-like lipids in animals. In insects, prostaglandins have been implicated in reproduction, fluid secretion, and activation of the immune system [72], inducing prophenoloxidase, phagocytosis and hemocyte spreading. None of the five prostaglandins annotated in our study has been functionally characterized in honey bees. Besides eicosanoids, we identified a second group of host-derived metabolites induced by the microbiota. These are three derivatives of juvenile hormone III **(Figure S2E)**, two of which were among the most discriminatory product ions **(Figure 3**, *m/z* 283.191 and *m/z* 269.176). Juvenile hormone III plays an important role in regulating growth, development, and reproduction of insects. In adult honey bees, it controls the pace of the developmental maturation from young nurse bees to older forager bees [73]. This process is linked to nutrition [74] and could therefore be affected by metabolic activities of gut bacteria. Notably, juvenile hormone derivatives in the gut may have local functions distinct from those in the brain or hemolymph, as was shown for heteropteran linden bugs [75].

### Gut metabolic profiles of colonized bees and hive bees show substantial overlap

To assess how much of the total metabolic output can be identified in hive bees under natural conditions, we analyzed the gut metabolome of 10-day old hive bees that experienced social interactions, natural dietary sources, and the native gut microbiota. Although principal component analysis revealed that hive bees clustered separately **(Figure S1B)**, we found that 27 of the 28 most discriminatory substrate ions and 15 of the 22 most discriminatory product ions showed qualitatively the same changes in hive bees as in CL bees (**Dataset S1**, Welch’s t-test, BH adj. *P* ≤0.01). On the substrate side, this included most flavonoid ions, all nucleosides, quinate, citrate, as well as the ions annotated as ω-hydroxy acids and phenolamides from the outer pollen wall. On the product side, we found four of the five prostaglandins and one of the juvenile hormone derivatives to be significantly increased in hive bees relative to MF bees suggesting that these host-derived metabolites are also induced under natural conditions. Moreover, ions corresponding to fermentation products were either significantly increased (sebacic acid, valerate) or showed a trend towards increased levels (succinate, pimelate) in hive bees. The same was the case for the four ions corresponding to possible degradation products of flavonoids (hydroxy‐ and dihydroxyphenylpropionate, maleylacetate, hydroxy-3‐ oxoadipate, **Dataset S1**). Overall, the remarkable overlap of metabolic changes between hive and CL bees highlights the relevance of our findings.

### Mono-colonizations explain 80% of the overall metabolic output of the honey bee gut microbiota

We thus far presented evidence for substrates and products of the complete microbiota in the honey bee gut. To elucidate which community members might be responsible for these transformations, we conducted mono-colonizations of MF bees with all seven bacterial species (again using four and two strains together for Firm-5 and *G. apicola*, respectively). All species successfully established in the gut of MF bees without other community members being present **(Figure S5)**. We again extracted metabolites from the mid-and hindgut of individual bees to address how many of the 372 robust ion changes can be explained by one or multiple mono-colonizations **(Dataset S1)**, i.e. show qualitatively the same change as in CL bees, using analysis of variance (ANOVA followed by Tukey HSD *post hoc* test at 99% confidence, *P* ≤0.05) **(Dataset S6–S7)**.

Remarkably, using these significance cutoffs 299 of the 372 (80%) robust changes between MF and CL bees could be explained by one or multiple mono¬colonizations. This included 201 (84%) substrate and 98 (74.%) product ions. The two *Lactobacilli* groups (Firm-5 and Firm-4) explained most changes, followed by *B. asteroides* and the two Gammaproteobacteria **(Figure 4A)**. Interestingly, the relative contribution to substrate conversion and product accumulation varied between mono-colonizations. For example, *B. asteroides* contributed relatively little to the conversion of substrates, but seemed to be responsible for the production of a relatively large fraction of bacterial products. The Firm-4 species showed the opposite pattern explaining relatively many bacterial substrates but a small fraction of bacterial products.

**Figure 4.**
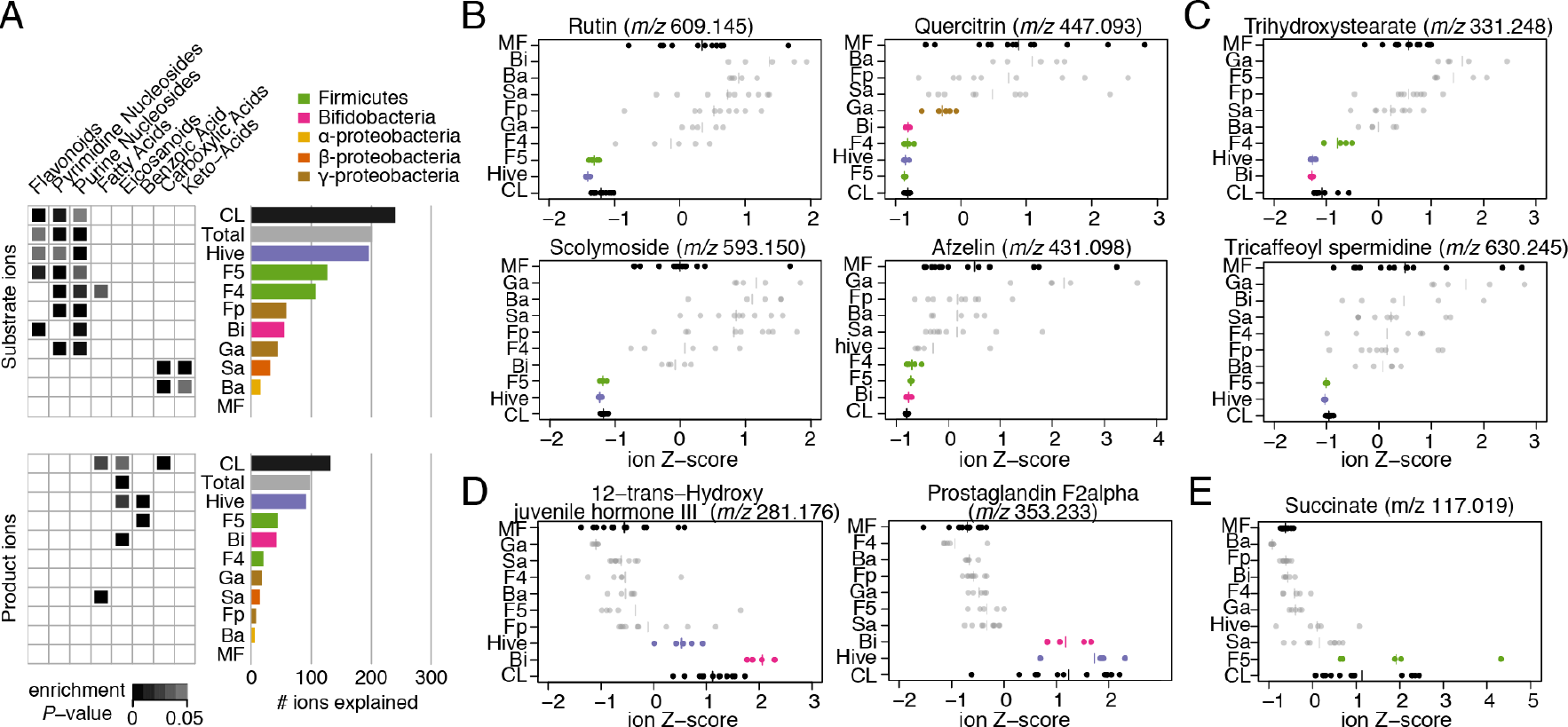
Overview of metabolite changes explained by different community members of the bee gut microbiota. (A) Distribution of metabolic changes explained by mono-colonizations. Bar graphs show distribution of the substrates (240 ions) and products (132 ions) separately. Enrichment P-values (one-sided Fisher's exact test) are provided for compound categories enriched in one or several mono-colonizations. The results of the full enrichment analysis are provided in **Dataset S2.** (B-E) Z-score transformed ion intensity of selected substrate and product ions are shown for all treatment groups. (B) glycosylated flavonoid substrates, (C) two substrates from the outer pollen wall, (D) two products corresponding to host-derived metabolites, (E) succinate, one of the major fermentation products. Groups depicted in color highlight treatment groups displaying a significant difference compared to MF bees in the same direction as the MF vs CL difference (one-way ANOVA, Tukey HSD *posthoc* test at 99% confidence, *P* ≤0.05). Plots for all 372 ions are provided in **Dataset S7.** MF, microbiota-free, Ba, *Bartonella apis* mono-colonized, Bi, *Bifidobacterium asteroides* mono-colonized, F4, Firm-4 mono-colonized, F5, Firm-5 mono¬colonized, Fp, *Frischella perrara* mono-colonized, Ga, *Gilliamella apicola* mono-colonized, Sa, *Snodgrassella alvi* mono-colonized, CL, colonized with the reconstituted microbiota, hive, hive bees.

#### Substrates explained by mono-colonizations

We again used enrichment analysis (one-sided Fisher’s exact test, P≤0.05), to get a high-level view of the functions of distinct community members in the conversion or production of certain compound classes **(Figure 4A)**. This revealed that all community members, except for *S. alvi* and *B. apis*, contributed to the disappearance of nucleosides in the bee gut (see also **Figure S2A)**. Remarkably, *S. alvi* and *B. apis* encode complete nucleoside biosynthesis pathways and therefore do not have to rely on external nucleoside resources. In turn, they seem to preferentially convert carboxylic acids and keto-acids in the gut (malate, *m/z* 133.0139, fumarate, *m/z* 115.003, citrate, *m/z* 191.019, a-ketoglutarate, *m/z* 145.014), which is consistent with the presence of complete TCA cycles and several carboxylate transporters in their genomes [32,54]. Flavonoids were enriched substrates for Firm-5 (11/126 substrate ions, *P*<0.01) and *B. asteroides* (6/55, *P*<0.01), and many flavonoids were also utilized by Firm-4 (7/107, P=0.051) **(Figure 4A, Figure S2B)**. Interestingly, rutin (*m/z* 609.145) and scolymoside (*m/z* 593.150) were exclusively depleted in the Firm-5 mono-colonization, while afzelin (*m/z* 431.098) was also utilized by Firm-4 and *B. asteroides*, and quercitrin (*m/z* 447.093) even by *G. apicola* **(Figure 4B)**. Similar patterns were also found for other flavonoids **(Figure S2B)** suggesting substrate specificity for the utilization of these pollen-derived compounds among community members.

In total, 27 of the 28 most discriminatory substrate **(Figure 3)** could be explained by at least one mono-colonization. The two ω-hydroxy acids ions from the outer pollen wall were exclusively utilized by *B. asteroides* and Firm-4 while the three phenolamides from the pollen coat were only depleted in the presence of Firm-5 **(Figure 4C, Figure S2C)**. In contrast, quinate and citrate were utilized by several community members suggesting that their almost complete depletion in CL and hive bees could be the result of a communal effort **(Dataset S1)**.

#### Products explained by monocolonizations

For bacterial products, 21 of the 23 most discriminatory ions for CL vs MF bees could be explained by at least one mono-colonization. Strikingly, *B. asteroides* explained the accumulation of all host-derived prostaglandins and was also responsible for the induction of two of the three juvenile hormone derivatives, suggesting that this community member has a distinct influence on the host **(Figure 4D, Figure S2E)**. Moreover, three major fermentation products that accumulated in CL bees could be explained by mono-colonizations. Succinate **(Figure 3E)** and pimelate ions were produced exclusively in bees colonized with Firm-5 and valerate only in the *B. asteroides* colonized bees **(Figure S2D, Dataset S1)**. Finally, we found that ions corresponding to putative flavonoid degradation products accumulated in bees colonized with Firm-4 and Firm-5 **(Figure S2B)**, which also explained most flavonoid utilization of the bee gut microbiota. This suggests that Firm-4 and Firm-5 do not only deglycosylate flavonoids, via expression of glycoside hydrolases [44], but may additionally degrade the aromatic backbone.

### *In vitro* recapitulation of metabolic functions of community members

Our *in vivo* results strongly suggest that specific gut bacteria utilize distinct substrates from the pollen diet of bees. This prompted us to test (i) whether the bacterial species could grow *in vitro* on a pollen-based culture medium and (ii) whether this would result in the metabolic conversions of the same compounds as was observed *in vivo.* To this end, we water-extracted metabolites from the same pollen batch that was used as a dietary source, and analyzed the metabolic composition of this extract using untargeted and targeted metabolomics. Detailed results are presented in the **Supplemental results 4** and show that pollen extracts contain physiologically meaningful levels of nutrients and are expectedly enriched in *amino acids and derivatives, flavonoids, monosaccharides,* and *carboxylic acids and derivatives* **(Figure S6)**. Strikingly, in the presence of this pollen extract, all community members, except for *S. alvi* displayed substantial growth compared to carbon-depleted or reduced control media in which little or no growth was observed after 16 h of incubation **(Figure 5A)**.

**Figure 5.**
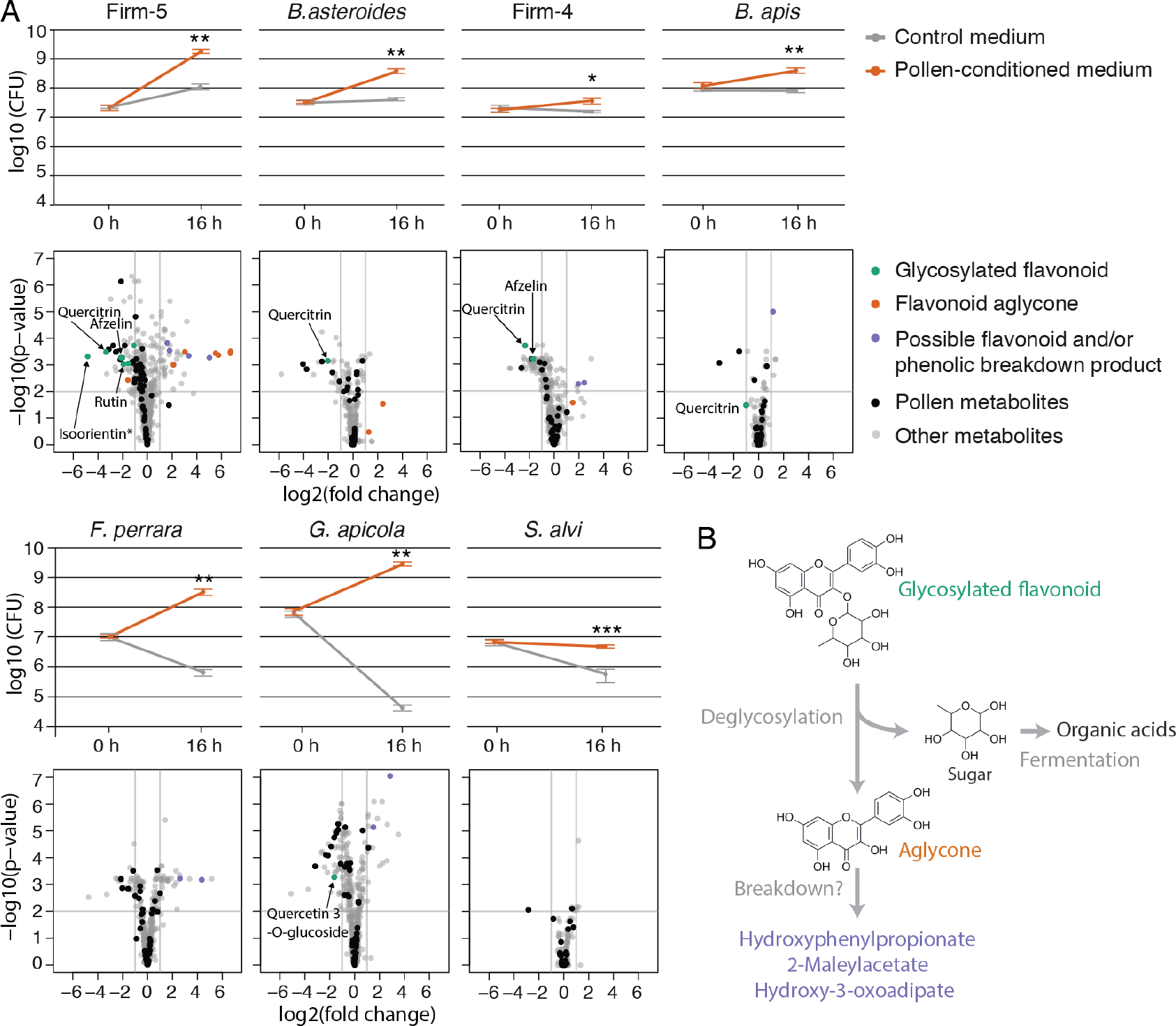
Recapitulation of metabolic changes induced by gut bacteria during *in vitro* growth in pollen-conditioned medium. (A) Line graphs show growth of each community member in control medium and pollen-conditioned medium based on colony-forming unit (CFU) counts at time-point 0 h and 16 h. Values are the mean of five replicates with error bars indicating standard deviation. n.s = not significant, * = p < 0.01, ** = p < 0.001, p < 0.0001 = *** (Welch t-test). Volcano plots of significance (Welch t-test BH adj. P-value) vs. log2(fold change) show metabolic changes in pollen-conditioned medium at time-point 16 h relative to 0 h. Ions identified as pollen-derived are highlighted in black. Ions annotated as glycosylated flavonoids, flavonoid aglycones (non¬glycosylated flavonoids), or putative flavonoid breakdown products are shown in color when they displayed log2(fold changes) >|1|. Other annotated ions are plotted in grey. (B) Model for the metabolism of flavonoids in the bee gut. Flavonoids are deglycosylated by specific bee gut bacteria resulting in the release of flavonoid aglycones. The sugar residues are likely fermented into organic acids. Accumulation of several (poly)phenol breakdown products *in vivo* and *in vitro* suggests that the aglycone may be broken down further.

We then profiled the metabolome in growth media before and after bacterial incubation in a separate metabolomics experiment. We annotated a total of 1031 ions **(Dataset S8)** of which 427 (41%) were also present among the 1079 ions from the *in vivo* dataset. In line with their growth profiles, the largest number of depleted metabolites (log2(FC) ≥|1| and Welch's t-test BH adj. *P* ≤0.01) were found for the growth cultures of Firm-5, followed by *G. apicola,* Firm-4, *F. perrara, B. asteroides, B. apis*, and *S. alvi* **(Figure 5A)**.

Using strict criteria we identified 17 ions - 13 pollen-derived substrates and 4 bacterial products – which were explained *in vivo* and *in vitro* by the same species **(Figure S7, Table S2)**. Seven of these 13 substrates belonged to the most discriminatory substrate ions for MF vs CL bees **(Figure 3)**: three flavonoids (quercitrin, afzelin, and rutin), one nucleoside (inosine) and ions corresponding to quinate, citrate, and 2-fuorate. The fact that multiple community members were responsible for the conversion of some of these substrates (*B. asteroides,* Firm-4, Firm-5, *F. perrara, B. apis* and *G. apicola*) demonstrates that our *in vitro* cultures allowed us to recapitulate metabolic activities covering the whole community.

We found remarkably overlapping substrate specificity for four flavonoids *in vitro* and *in vivo* with the Firm-5 species being the only member capable of converting rutin and scolymoside, while quercitrin and afzelin were also utilized by Firm-4, and quercitrin additionally also by *B. asteroides* and *B. apis* **(Figure 5A)**. Among the four *in vitro* recapitulated products were three of the four ions corresponding to putative breakdown products of flavonoids **(Figure S7, Table S2)**. These ions accumulated *in vivo* and *in vitro* in the presence of Firm-4 and/or Firm-5 providing further evidence for breakdown of the polyphenolic ring structure of flavonoids. However, we also found that deglycosylated flavonoids (i.e. aglycones) accumulated in cultures of Firm-4, Firm-5 and *B. asteroides* **(Figure 5A)**. Based on this data we suggest that flavonoid degradation involves two steps **(Figure 5B)**: (i) deglycosylation of sugar residues and their subsequent fermentation and (ii) the breakdown of the polyphenol backbone. The second step could be relatively slow explaining why aglycones accumulated *in vitro* (16 h), but not *in vivo* (10 days).

An obvious difference in our *in vitro* experiments compared to the *in vivo* situation is the presence of the host, which may pre-digest pollen grains before gut bacteria utilize pollen-derived metabolites. For example, certain sugars and amino acids are expected to be present in low amounts *in vivo* because of host absorption. Conversely, the host may also provide metabolites that support growth of some community members. This could explain the poor growth of *S. alvi in vitro*, especially since *in vivo S. alvi* is tightly associated with the gut epithelium and other gut bacteria such as *G. apicola* [76].

### Evidence for cross-feeding in the bee gut mocrobiota

Microbial species in gut communities can organize into food chains, where one species provides metabolites that can be utilized by others. Such metabolites may be released from insoluble dietary particles via bacterial degradation or can be generated as waste products of metabolism [2]. To identify possible metabolic interactions between community members of the bee gut microbiota, we focused on ions that *in vivo* significantly accumulated in some mono-colonizations and were depleted in others **(Dataset S1)**. A total of 27 ions showed such opposing changes between two or several mono-colonizations **(Table S3)**.

A striking example of a potential metabolic food chain was the liberation and consumption of one of the major bacterial substrates in CL bees, 9-10-18‐ trihydroxystearate (*m/z* 331.248), originating from the outer pollen wall. In our mono-colonization experiments, the corresponding ion was depleted in Firm-4 and *B. asteroides*, but accumulated in the case of Firm-5 and *G. apicola* **(Figure 4C)**. This suggests that the latter two species facilitate the release of this w‐ hydroxy acid from the outer pollen wall making it more accessible for degradation by Firm-4 and *B. asteroides.*

A second example is the ion annotated as pyruvate (*m/z* 87.008), which substantially accumulated in the gut of bees mono-colonized with *G. apicola*, but was utilized as a substrate by other bacteria such as *S. alvi* and Firm-5 **(Figure 6A)**. A syntrophic interaction between *G. apicola* and *S. alvi* had previously been suggested, because they are co-localized on the epithelial surface of the ileum [76], and harbor complementary metabolic capabilities [32]. To test for potential cross-feeding of pyruvate from *G. apicola* to *S. alvi*, we supplemented the pollen extract-based medium of *S. alvi* with spent supernatant of *G. apicola* cultures. While growth of *S. alvi* was only weakly improved compared to the control medium **(Figure S8)**, metabolome analysis **(Dataset S9)** confirmed that pyruvate and five other ions **(Figure 6B)**, which accumulated during the growth of *G. apicola*, were utilized from the conditioned medium by *S. alvi.* The other ions included three fermentation products, a nucleoside derivative, and hydroxyphenylpropionate, which could be a breakdown product of flavonoid degradation by *G. apicola.* These results confirm our predictions from the *in vivo* dataset and show that bee gut bacteria engage in cross-feeding interactions among each other. While not essential for gut colonization in itself as based on our mono-colonization experiments, such interactions are likely important for community resilience and reflect the longstanding coexistence among community members.

**Figure 6.**
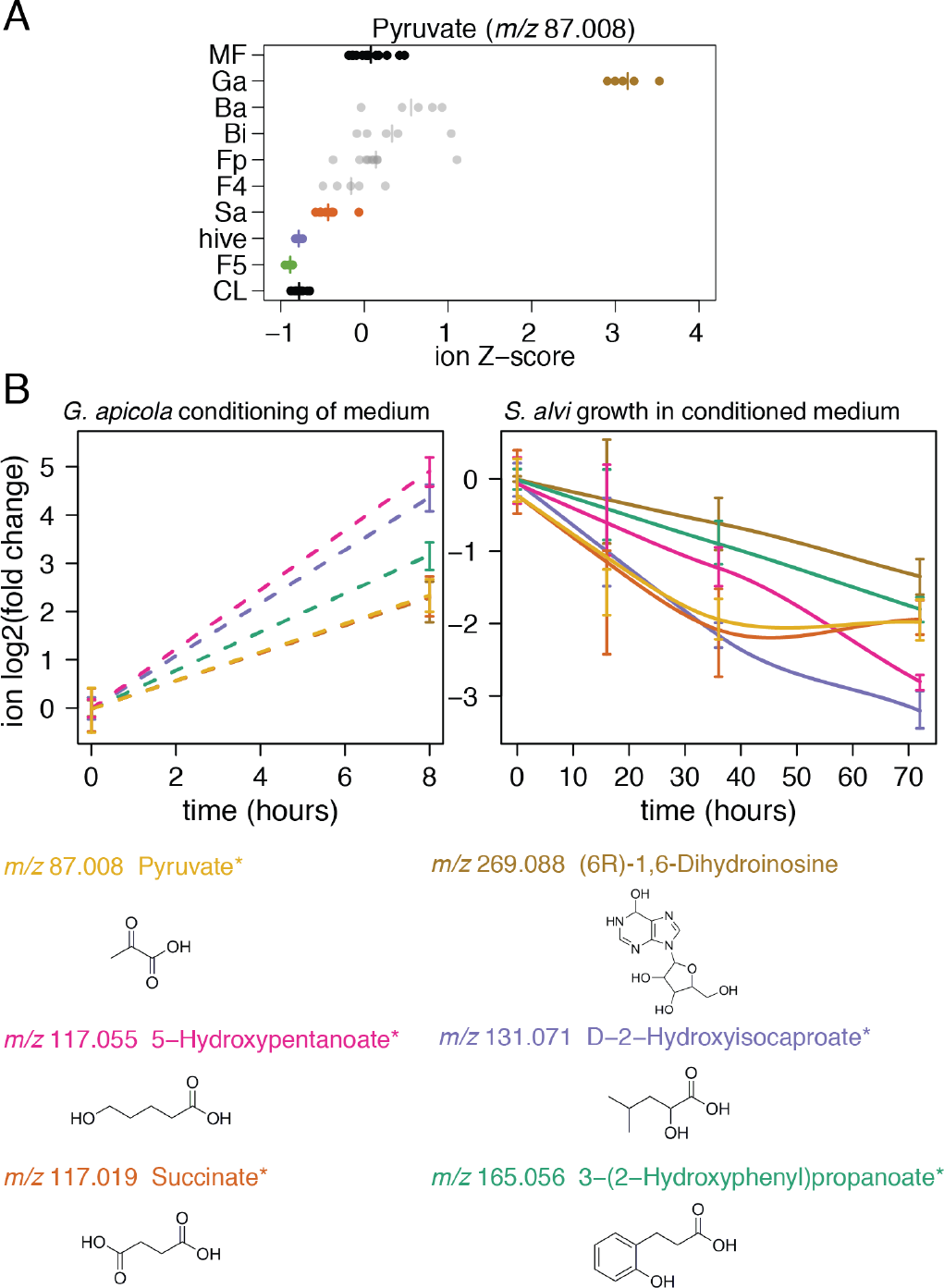
Cross-feeding between *G. apicola* and *S. alvi.* (A) Evidence for cross¬feeding of pyruvate in the honey bee gut. Z-score transformed ion intensities reveal that the ion annotated as pyruvate accumulates in bees mono-colonized with *G. apicola*, but is depleted in hive bees, CL bees, and bees mono-colonized with *S. alvi* and Firm-5. (B) Six potentially cross-fed ions which accumulated during *in vitro* growth of *G. apicola* (left sub-panel) and were consumed by *S. alvi* when it was grown on the *G.* apicola-conditioned medium mixed 1:1 with fresh medium (right sub-panel). Smoothed lines are added for interpretation purposes only and are dashed in the left panel because they are drawn through two points only. Error bars represent the standard deviation based on three replicate cultures. Chemical structures of the first annotation of each ion are shown. Asterisks indicate ions with ambiguous annotations.

## Conclusions

The simple composition and experimental amenability of the honey bee gut microbiota facilitated our systems-level approach resulting in the characterization and disentangling of the metabolic potential of a complete gut microbial ecosystem. We reconstituted the honey bee gut microbiota from cultured strains, characterized the metabolic output of the complete microbiota, identified the contributions of individual community members *in vivo*, and recapitulate their activities *in vitro.* Our results provide unprecedented insights into the metabolic functions of bee gut bacteria.

As in the mammalian and termite gut ecosystem [1,67], we conclude that most substrates utilized by the bee gut microbiota are indigestible compounds originating from the hosts’ diet **(Figure 7)**. These include plant metabolites from the outer pollen wall, such as ω-hydroxy acids, phenolamides and flavonoid glycosides. While one of the bee gut bacteria had previously been identified to utilize a major pollen polysaccharide (pectin), our data provides the first evidence for a role of the gut microbiota in breaking down outer pollen wall components. Bacterial fermentation of these pollen-derived compounds resulted in the accumulation of organic acids (e.g. succinate) and polyphenol degradation products, which are likely to impact the physicochemical conditions in the colonized gut. In addition, we found that host-derived signaling molecules are induced by *B. asteroides*, suggesting a specific interaction of this gut symbiont with the host.

**Figure 7.**
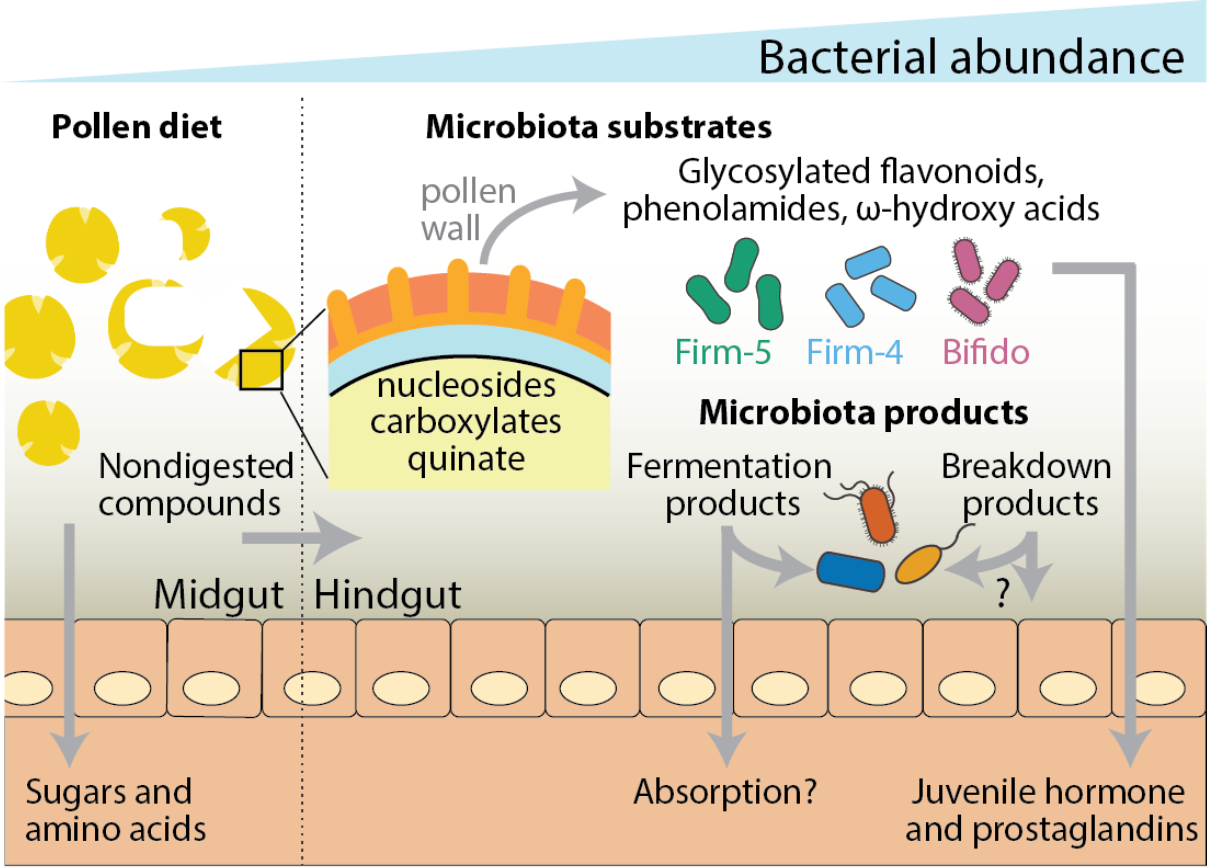
A summary of the metabolic activities of the bee gut microbiota identified in this study. Pollen is likely predigested in the midgut where bacterial levels are relatively low [45]. Here, the host absorbs accessible pollen‐ derived compounds such as simple sugars (glucose or fructose) and amino acids [46,47]. Non-digested pollen grains and pollen compounds enter the hindgut where bacterial density is much higher. We found nucleosides, various carboxylic acids (e.g. citrate, malate, fumarate) and aromatic compounds (such as quinate) from pollen to be utilized by bee gut bacteria. In the posterior part of the hindgut, three community members (Firm-5, Firm-4, and *B. asteroides)* metabolize major components of the outer pollen wall, including flavonoids, phenolamides and ω-hydroxy acids. The metabolic activities of the microbiota lead to the accumulation of fermentation products and intermediates of aromatic compound degradation. Some of the bacterial products may be utilized by other community members, as exemplified by the cross-feeding between *G. apicola* and *S. alvi*, or absorbed by the host. In addition, the gut symbiont *B. asteroides* seems to increase the production of several host metabolites (juvenile hormone derivatives and prostaglandins) that have key functions in immunity and physiology.

Based on the mono-colonization experiments, we further conclude that most metabolic output of the bee gut microbiota can be explained by activities of individual community members. These bacteria seem to occupy largely independent yet overlapping metabolic niches in the gut. While we have found evidence for cross-feeding interactions (e.g. between *G. apicola* and *S. alvi*), our findings generally indicate that coevolution of gut bacteria does not necessarily result in interdependencies. This may be generally the case for gut bacteria as they cannot rely on the presence of specific interaction partners in the highly dynamic gut environment, but rather adapt to diet-derived nutrients.

The metabolic activities identified in this study are likely key for the symbiotic functions of the bee gut microbiota and thus may be directly linked to its impact on bee health and physiology [34,41]. The fact that we identified many of these metabolic changes in hive bees under natural conditions demonstrates the relevance of our findings and validates the gnotobiotic bee model. Moreover, our study highlights the versatility of high-throughput untargeted metabolomics to disentangle metabolic functions in microbial ecosystems. We believe that this systematic approach can be extended to other gnotobiotic animals to enable a better understanding of the diversity of metabolic activities and functions that are present in microbial communities.

## Materials and Methods

### Honey bee experiments

MF bees were obtained from a healthy looking colony of *Apis mellifera carnica* located at the University of Lausanne. To this end, dark-eyed pupae were carefully removed from capped brood cells with sterile tweezers and transferred to sterilized plastic boxes as described previously [36]. Boxes with pupae were kept with a source of sterile sugar water (50 % sucrose solution, w/v) at 35°C with 80 – 90 % humidity for the first two days. The temperature was reduced to 32°C on the third day. One or two bees per box were dissected one day prior to colonization and their homogenized hindguts (in 1 ml 1x PBS) were cultured on media as described below to check for sterility of the bee gut. Only healthy looking emerged bees without any deformations and sterile guts were selected for experiments. For colonization of MF bees, bacterial strains were inoculated from glycerol stocks and restreaked twice. Details on bacterial strains and culture conditions can be found in **Table S1**. Bacterial cells were harvested and resuspended in 1x PBS/sugar water (1:1, v/v) at an OD600 of 1. For colonization, bacterial suspensions were added to a source of sterilized pollen and provided to the MF bees (for details see **Supplemental methods**). The mid-and hindgut of gnotobiotic bees were dissected at day 10 post colonization and stored at -80°C until further use. To obtain age controlled hive bees, several brood frames without adult bees were transferred from the hive to a ventilated Styrofoam box kept in an incubator at 32 – 34°C in the laboratory overnight. The next day, newly emerged bees were collected, marked on the thorax with a pen and reintroduced into the hive. These bees were recollected 10 days later, and their mid-and hindguts were dissected and stored at -80°C until further use. This experiment was repeated at two different time-points of the year (spring and fall, referred to as experiment 1 and experiment 2 in this study). Whenever possible, we included bees from both experiments in our analysis, such as for CL and MF bees. However, this was not possible for all mono-colonizations due to bacterial contaminations (as detected by qPCR) or in a few cases due to the presence of above-threshold viral loads. The precise numbers of bees included per condition are listed in **Table S4**.

### Determining bacterial colonization levels in honey bees

Colonization levels of gut bacteria were determined by qPCR using species‐ specific primers on DNA samples obtained from the gut tissues also used for metabolomics analysis. Details on DNA/RNA extraction methods are given in **Supplemental methods.** Each DNA sample was screened with eleven different primer pairs targeting the actin gene of *Apis mellifera*, the 16S rRNA gene of all bacteria, and the 16S rRNA gene of nine specific bacterial species (the seven species used this study and two non-core species frequently found in the gut of *Apis mellifera*, Alpha-2.1 and *Lactobacillus kunkeei).* Primers used for this qPCR analysis are summarized in **Table S5.** We also screened all gut samples for the presence of viruses. Samples that were contaminated with other bacteria than the desired ones or which had high virus titers were excluded from the analysis. The MIQE guidelines (minimum information for publication of qPCR experiments) were followed throughout the data analysis of the qPCR experiments [77]. Details on the qPCR analysis can be found in **Supplemental methods.**

### *In vitro* growth on pollen extracts

Bacteria were pre-cultured on solid media from -80°C glycerol stocks before liquid cultures were inoculated for *in vitro* growth experiments. For *G. apicola* ELS0169, *S. alvi* wkB2, *F. perrara* PEB0191, and *B. apis* PEB0149, we used a modified M9 minimal medium supplemented with casamino acids and vitamins. For *B. asteroides* ELS0168, the Firm-5 strains, and Firm-4 Hon2N^T^, we used carbohydrate-free MRS (cfMRS) medium [78]. Bacteria were harvested from plates or spun down from overnight liquid cultures (the latter only for *Lactobacilli* and *B. asteroides*) and resuspended in the corresponding minimal medium. Freshly prepared liquid cultures were supplemented with either 10% (v/v) ddH_2_O or pollen extract and inoculated at a final OD600 of 0.05 (see **Supplemental methods** for details on pollen extract preparation). Half of the culture was immediately processed to determine colony-forming units (CFUs) and to harvest supernatants for metabolomics at time point 0 h, i.e. before growth. The other half of the culture was incubated for 16 h according to the conditions listed in **Table S1** and then processed in the same way. For CFU counting, serial dilutions were plated on solid media and incubated under the species-specific culturing conditions. For metabolomics analysis, the remaining bacterial culture was spun down at 20,000x g at room temperature for 10 min and 300 [il of spent culture supernatant transferred to a fresh tube stored at – 80°C until further processing. Five replicates were included for each species and treatment group.

For the cross-feeding experiment between *G. apicola* and *S.alvi, G.apicola* strain ESL0169 was grown for 8 h in pollen-supplemented M9 medium as described before. Cultures were sterile-filtered and mixed with fresh pollen-supplemented M9 medium 1:1 (v/v). For the control condition, non-inoculated cultures were incubated for 8 h, sterile filtered and mixed with fresh pollen‐ supplemented M9 medium 1:1 (v/v). Then, *S. alvi* was added to each tube at a final OD_600_ of 0.05. Growth of *S. alvi* was assessed by OD600 measurements. For metabolomics analysis, supernatants were sampled at time points 0 h and 8 h for the *G. apicola* conditioning cultures and at time points 0, 16, 36, 72 h for the *S. alvi* cultures in conditioned medium.

### Metabolite extraction and profiling

Metabolites from gut and pollen samples were water-extracted after mechanical disruption, and supernatants from the *in vitro* experiments were harvested by centrifugation. Detailed methods can be found in the **Supplemental methods.** Gut samples were pre-selected based on their wet-weight (arithmetic mean 55.1 mg, standard deviation 9.9). Ten times more water than the gut wet-weight (v/w) was added, and the samples were homogenized with 0.1 mm zirconia beads in a Fast-Prep24^TM^5G homogenizer at 6 m/s for 45 s. This homogenate was snap-frozen in liquid nitrogen for subsequent DNA/RNA extraction, and 20 times diluted for metabolite extractions which was achieved by incubating the samples in a pre-heated thermo-mixer at 80°C and 1400 rpm for 3 min. After each minute, the samples were vortexed for 10 s. Subsequently, the samples were centrifuged at 20000x g and 4°C for 5 min, and 150 [il of the resulting supernatant was transferred to a new tube and centrifuged again. Finally, the supernatant was diluted 10x and stored at -80°C or on dry ice for subsequent metabolomics analysis.

Samples were injected into an Agilent 6550 time-of-flight mass spectrometer (ESI-iFunnel Q-TOF, Agilent Technologies) operated in negative mode, at 4 Ghz, high resolution, and with a mass / charge (*m/z*) range of 50-1000 [21]. The mobile phase was 60:40 isopropanol:water (v/v) and 1 mM NH4F at pH 9.0 supplemented with hexakis (1H, 1H, 3H‐ tetrafluoropropoxy)phosphazine and 3-amino-1-propanesulfonic acid for online mass correction. After processing of raw data as described in [21], *m/z* features (ions) were annotated by matching their accurate mass to sum formulas of compounds in the Kyoto Encyclopedia of Genes and Genomes (KEGG) database with 0.001 Da mass accuracy and accounting for deprotonation [M-H+]\ This method cannot distinguish between isobaric compounds, e.g. metabolites having identical *m/z* values. The raw data of samples from the three sets of experiments (bee gut samples, *in vitro* supernatants, and cross-feeding supernatants) were processed and annotated separately to accommodate their different sample matrices or times of measurement. This data can be explored in **Dataset S1, S8, and S9.** Raw data processing and annotation took place in Matlab (The Mathworks, Natick) as described previously [21] and downstream processing and statistical tests were performed in R (R Foundation for Statistical Computing, Vienna, Austria).

Selected metabolite samples were measured in targeted fashion using ultra-high-pressure chromatography-coupled tandem mass spectrometry as described before [71]. Metabolite quantifications were performed by interpolating observed intensities to a standard curve of the metabolite.

Flavonoid ions were targeted for MS/MS fragmentation as [M-H^+^] electrospray derivatives with a window size of ± 4 m/z in Q1. Fragmentation of the precursor ion was performed by collision-induced dissociation at 0, 10, 20, and 40 eV collision energy and fragment-ion spectra were recorded in scanning mode by high-resolution time-of-flight MS. Spectra were interpreted using MetFrag (Ruttkies *et al.*, 2016), and spectral cosine similarity scores were calculated between reference spectra that were obtained in-house or library spectra from MassBank of North America (MoNA, http://mona.fiehnlab.ucdavis.edu/). For further details see **Supplemental methods.**

### Untargeted metabolomics data analysis

All steps of the downstream data analysis were performed in R (R Foundation for Statistical Computing, Vienna, Austria). Samples from double injections (technical replicates) were confirmed to be highly similar and averaged. Subsequent analyses were performed on these averaged ion intensities, which are available in **Dataset S1, Dataset S8, and Dataset 9**.

Ions that were deemed robustly different between the MF and CL bees were those that were significantly different (Welch t-test, BH adj. *P* ≤0.01) between CL and MF in both independent experiments. Differences between MF and CL samples were expressed as log2(fold-change) values for both experiments separately and for pooled data of both experiments (see **Dataset S1**).

Enrichment analyses were computed on compound class categories from KEGG (in house database), which are added in the column “compound class” in **Dataset S1**. Some ions with ambiguous annotations had a compound class associated to multiple of these annotations. However, supported by the observation that compound classes between alternative annotations were often the same (or highly related) only the compound class of the first annotation was used as input for one-sided Fischer’s exact tests **(Dataset S3)**.

One-way analysis of variance (ANOVA) was performed between all bee gut samples after selecting the relevant samples from the data matrix and normalizing the intensities to ion standard (Z-) scores (i.e. by row). The results of the full ANOVA analysis can be explored in **Dataset S6**. For this study the focus was on differences between any group and MF bees which were considered significant when having an Tukey HSD *post hoc* adjusted P-value 0.05. When for a specific monocolonization group this significance cut-off was met and the direction of the change was the same as that for CL vs MF, the ion was considered to be “explained” by this group.

In order to enrich for pollen ions we only considered ions with an arithmetic mean intensity of ≥10,000 in the pollen samples, in addition to being highly significantly different from water-matrix control samples (Welch t-test, BH adj. ≤ 0.001) and displaying a large (> 2) log2(fold change) difference.

For the *in vitro* data (Dataset S8), the goal was to identify pollen substrates and bacterial products for which changes in levels were observed *in vivo* and *in vitro.* Pollen ions were mapped by matching the top annotation formula of both datasets. For all media-strain combinations we performed a statistical comparison between time point 16 h and 0 h and considered only those ions with a log2(fold change) of ≥|1| and BH adj. P-value of ≤0.01 as significant *in vitro* products or substrates. In order to be certain that only pollen‐ derived substrates were included, for every strain only ions that displayed a significant negative log2(fold change) exclusively in the base medium supplemented with pollen extract were considered as *in vitro* pollen substrates.

To identify ions that might be cross-fed between *G. apicola* and *S. alvi,* ions were selected that increased during the growth of *G. apicola* and were depleted when *S. alvi* was grown in this conditioned medium mixed 1:1 with fresh base medium. To do this all ion intensities for both strains **(Dataset S9)** were split and transformed to log2(fold-change) with respect to the first time point of sampling. Ions that had a total log2(fold change) of ≥|1| during *G. apicola* growth and a total log2(fold change) of ≤|−1| during *S. alvi* growth were selected.

For a more extensive version of these methods including settings of functions, and detailed protocols see Supplemental methods.

## Author Contributions

Conceptualization, L.K., R.A.T.M., K.E., U.S., and P.E.; Methodology, L.K., R.A.T.M., and P.E.; Validation, L.K., R.A.T.M., and P.E.; Investigation, L.K., R.A.T.M., and P.E.; Supervision, U.S. and P.E.; Writing – Original Draft, L.K., R.A.T.M., and P.E.; Funding Acquisition, U.S. and P.E.

## Acknowledgements

The authors would like to acknowledge Dr. Emma Schymanski for excellent help with the interpretation of MS/MS fragmentation spectra. This work was supported by the University of Lausanne and the ETH Zürich, the SNSF research fund 31003A_160345 (to P.E.), and the ERC Starting grant 714804 – MicroBeeOme (to P.E.). R.A.T.M. was supported by the ERA-Net for Applied Systems Biology ERASysAPP (to U.S.).

